# Synthetic Generation of Dynamic Omics Data Demonstrates *Aspergillus nidulans* BrlA Paradoxical Wall Stress Response

**DOI:** 10.1101/2025.03.02.638868

**Authors:** Joseph Zavorskas, Harley Edwards, Walker Huso, Alexander G. Doan, Mark R. Marten, Steven Harris, Ranjan Srivastava

## Abstract

We propose a method to generate additional dynamic omics trajectories which could support pathway analysis methods such as enrichment analysis, genetic programming, and machine learning. Using long short-term memory neural networks, we can effectively predict an organism’s dynamic response to a stimulus based on an initial dataset with relatively few samples. We present both an *in silico* proof of principle, based on a model that simulates viral propagation, and an *in vitro* case study, tracking the dynamics of *Aspergillus nidulans’* BrlA transcript in response to antifungal agent micafungin. Our *silico* experiment was conducted using a highly noisy dataset with only 25 replicates. This proof of principle shows that this method can operate on biological datasets, which often have high variance and few replicates. Our *in silico* validation achieved a maximum R^2^ value of approximately 0.95 on our highly noisy, stochastically simulated data. Our *in vitro* validation achieves an R^2^ of 0.71. As with any machine learning application, this method will work better with more data; however, both of our applications attain acceptable validation metrics with very few biological replicates. The *in vitro* experiments also revealed a novel paradoxical dose-response effect: transcriptional upregulation by *Aspergillus nidulans* BrlA is highest at an intermediate dose of 10 ng/mL and is reduced at both higher and lower concentrations of micafungin.

## Introduction

Multi-omics methods amalgamate multiple sources of biological “big data,” each of which often describe the organism-wide dynamics of some part of the central dogma of biology: DNA -> RNA -> protein.^1^ For example, a multi-omics study might consider a regulatory network containing transcription factors, kinases/phosphatases, repressor proteins, etc. all responding to a certain stimulus. To collect transcriptomic data, for example, one could use RNA sequencing (RNA-seq) to understand the dynamics of all transcripts in an organism.^2^ Alternatively, DNA microarrays^3^ or quantitative polymerase chain reaction (qPCR)^4^ can be used to study the dynamics of a small subset of transcripts. To perform studies from a protein lens, one can use mass-spectrometry based proteomics to collect data on all proteins present in the cell at the time of sampling.^5^ This analysis can be extended to understand protein regulation via post translational modifications by searching for additions such as phosphorylation, methylation, etc.^6^

Interactions between genes and signaling pathways are both complicated and interconnected. Commonly, researchers will select pathways of interest to focus their data analysis and reduce complexity. To glean connections without a pathway basis, techniques such as pathway enrichment analysis,^7,8^ genetic programming (GP),^9–11^ and machine learning (ML) can be used.^12,13^ Each of these data analysis methods benefits from greater quantities of quality data, especially GP and ML. However, many of the data collection methods mentioned previously require significant time and energy to generate quality replicates. In addition, several time points are required to accurately capture the dynamics of an organism’s response to a stimulus, each of which requires both biological and technical replicates.^14^

This work seeks to generate synthetic, multi-omics replicates which could be used to further biological pathway understanding, such as GP algorithms or machine learning. We used a long short-term memory (LSTM) neural network^15^ to capture the dynamic response of *Aspergillus nidulans* BrlA to varying doses of the echinocandin antifungal agent micafungin.^16,17^ Before this case study, we conducted proof of concept studies for the LSTM on a viral propagation model.^18^ In this model system, significant noise was introduced via stochastic simulation of the differential equations, to simulate the high variance of true biological data.

### Long Short-Term Memory Neural Networks

To create mechanistic models for dynamic gene expression, it is more important that the technique used to generate synthetic -omics replicates consider the temporal dynamics between points than the individual points themselves. These requirements can be satisfied using machine learning techniques that perform sequence learning^19^ such as recurrent neural networks (RNNs), and their extension, LSTMs.^15^ RNNs are layered algorithms which process sequential information by taking one point as input in each layer and producing a single output.^20^ RNNs’ strength is their built-in memory, where hidden nodes of the current layer pass information to the hidden nodes of the subsequent layer.^21^ However, RNNs have a few fundamental flaws that make them unsuitable to apply to large problems with long time scales. Most importantly, RNNs face an issue known as the vanishing gradient problem.^22^ This problem is caused by the consistent scaling of information by each layer’s activation function by values between -1 and 1.^23^ Therefore, the longer the sequence an RNN is considering, the more extreme the vanishing gradient problem is expected to become.^24^ The exact opposite effect, exploding gradients, is also possible depending on the activation function and scaling type used.^25^

LSTMs are a subset of RNNs that were created by Hochreiter and Schmidhuber^15^ to address the shortcomings of traditional RNNs. A typical LSTM, as shown in **Figure 1**, contains two streams of data, the “cell” stream and the “hidden state” stream. The cell stream contains the long-term memory of the network, while the hidden state directly handles the input of the current time point. LSTMs typically contain three “gates” which apply activation functions to the data streams: the forget gate, input gate, and output gate. The forget gate decides what information from the cell memory stream to delete based on what it has learned is important to current and future time points and discards less needed information.^2627^ The input gate uses multiple activation functions to store the current hidden state information in the memory stream. Finally, the output gate combines information from memory and the current input to generate the output at the current time point. An activation function is never permanently applied to the memory stream, only to a copy of its value in the output gate, solving the vanishing gradient problem.

## Case Studies

### Viral Propagation

First, we validate our LSTM approach by using the simple biological model proposed by Srivastava *et al*. (2002), which represents the infection and propagation dynamics of a generalized virus.^18^ Their model simplifies the viral infection process into three general stages: **1)** *tem*, the free, translated genetic template responsible for creating the physical components of the virus; **2)** *gen*, the genomic nucleic acid sequence that is packaged within new viruses to infect other cells; **3)** *struct*, the structural proteins such as capsid proteins or envelope proteins that form the physical virus.^18^ Most importantly, *gen* produces *tem*, which can produce both *gen* and *struct*. A complete virus is created by the export of *gen* and *struct* together. This system can be represented by three coupled differential equations, as shown in **Equation 2**:

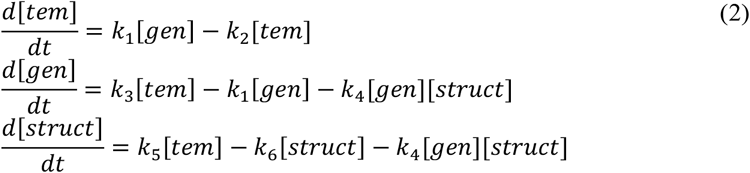

The parameters in these equations and the processes they represent are included in **Table 1**. This viral propagation model tests the LSTM’s ability to simulate a system with a dynamic phase which eventually leads to a stable steady state. This steady state occurs at *tem, gen*, and *struct* values of 20, 200, and 10000, respectively.^18^ Our goal for the LSTM is to generate accurate dynamic trajectories representing differing initial infection quantities by adjusting the initial condition for *tem*.

**Table 1.**
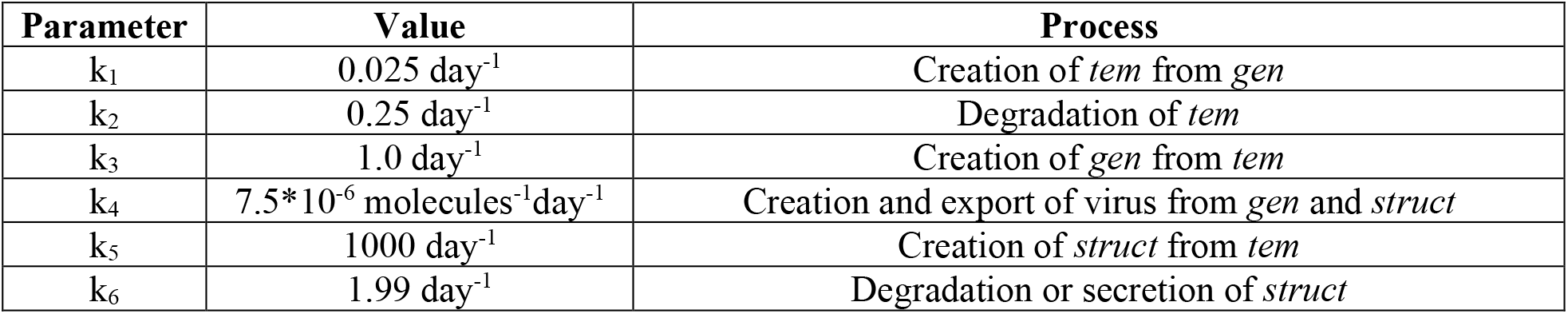
The rate constants for each viral propagation reaction are included below. The corresponding process for each consumption or production term is also included. Rate constants were chosen by Srivastava *et al*. (2002) to result in a stable steady state.

| Parameter | Value | Process |
| --- | --- | --- |
| $k_1$ | $0.025 \text{ day}^{-1}$ | Creation of *tem* from *gen* |
| $k_2$ | $0.25 \text{ day}^{-1}$ | Degradation of *tem* |
| $k_3$ | $1.0 \text{ day}^{-1}$ | Creation of *gen* from *tem* |
| $k_4$ | $7.5 \times 10^{-6} \text{ molecules}^{-1} \text{ day}^{-1}$ | Creation and export of virus from *gen* and *struct* |
| $k_5$ | $1000 \text{ day}^{-1}$ | Creation of *struct* from *tem* |
| $k_6$ | $1.99 \text{ day}^{-1}$ | Degradation or secretion of *struct* |

### Aspergillus nidulans BrlA

Various species from the genus *Aspergilli* are widespread in nature and are used prevalently in the bioprocess industry. For example, *A. oryzae* and *A. niger* are used for many bioproduction processes such as the fermentation of sake and soy sauce, production of citric and gluconic acids, and the production of many useful enzymes. ^28–30^ However, other *Aspergilli* are opportunistic human pathogens, the most virulent of which is *A. fumigatus*. This fungus can cause aspergillosis, especially in hosts which are already immunocompromised.^31^ *A. fumigatus* is so virulent due to its aggressive spore production and dissemination, which when breathed in are quickly cleared by the immune system.^32^ However, these ubiquitous spores can easily infect immunocompromised individuals who are frequently exposed to them in everyday life.

In *Aspergilli*, fungal spores are produced through an asexual reproduction process called conidiation. This process is regulated by a genetic pathway that responds to both light and air to decide when to initiate spore production.^33^ The signaling pathways that initiate conidiation all feed into and activate the central transcription factor of asexual reproduction, BrlA, which is the first of three central transcription factors in the asexual reproduction pipeline, organized as follows: *brlA*→ *abaA*→ *wetA*.^34^ Each transcription factor is responsible for a different phase of conidial development, which is evidenced by differing morphological defects in their respective deletion mutations.^35,36^ Understanding the role of BrlA, and the central conidiation pathway, is very important to understand fungal reproduction, aspergillosis, and other pathogenic fungi.

However, according to literature sources^37,38^ and our previous findings ^39,40^, BrlA appears to have functions outside of the asexual reproduction pathway. Notably, BrlA appears to play a role in cell wall stress response. This function is particularly significant because echinocandins, one of the most potent classes of antifungal agents, specifically target the fungal cell wall.^16,17^ Interestingly, in *Aspergillus* species, echinocandins produce only a fungistatic effect rather than a fungicidal one, suggesting these organisms possess inherent resistance mechanisms against these drugs.^41^

There is a significant amount of evidence that BrlA participates in the cell wall stress response. First, the *brlA* transcript was found to be significantly upregulated in response to micafungin exposure in our previous work.^39^ Additionally, this transcriptional upregulation occurred without the upregulation of the transcription factors *abaA* and *wetA*, which are commonly thought to act together.^36^ Second, the terminal MAP kinase of the cell wall integrity (CWI) signaling pathway, MpkA, has been shown to physically associate with, and have transcriptional effects, on BrlA. Kovacs *et al*. found that *brlA* transcription changed significantly in an *mpkA* deletion mutant.^38^ Rocha *et al*. found that MpkA and BrlA were “physically associated” during asexual reproduction via coimmunoprecipitation (Co-IP).^37^ Considering that MpkA is a serine/threonine kinase, this physical association may indicate that BrlA is being phosphorylated by MpkA to carry out cell wall integrity-related processes. A final piece of evidence supporting the hypothesis that BrlA participates in the cell wall stress response is that the transcriptional upregulation of *brlA* in response to micafungin does not occur in an *mpkA* deletion mutant.^42^

Our group has also explored possible downstream gene targets that *brlA* may regulate as part of the cell wall stress response we are proposing. Because of our interest in *brlA* transcriptional dynamics, we use it as an experimental case study in this paper. We collected detailed dynamic data of *brlA*’s transcriptional response to micafungin exposure for use in a proof-of-concept for generating synthetic dynamic -omics trajectories. Ultimately, these data and any synthetic trajectories could be used in a genetic programming approach to understand the upstream regulation of *brlA* in either a conidiation or cell-wall stress context. Future work may include computational analysis of *brlA*’s transcriptional network using this new tool.

## Methods

### Viral Propagation Proof of Principle

We generate a noisy, sparse solution to Srivastava *et al*.’s viral propagation model by generating a stochastically simulated dynamic trajectory and sampling it at evenly spaced time intervals. Biological data is often extremely noisy and does not have good time resolution due to the high cost of an individual biological replicate. To stress test the LSTM on more biologically realistic data, we introduced noise by performing stochastic simulation of the differential equations using the Gillespie stochastic simulation algorithm (SSA).^43^ In addition, we created sparse dynamic trajectories by removing all but a few small “sampling” events from the data. Sampling the generated noisy trajectories at evenly spaced time intervals

In a single step of the Gillespie simulation, the following occurs: **1)** the “propensity” of each reaction is calculated, essentially the likelihood that the reaction will occur given the current conditions; **2)** a random number is generated and subjected to the following equation to calculate the size of the current time-step. **3)** based on the calculated propensities, the first reaction to fire is calculated. Gillespie assumes that this reaction is the only one that fires in the short timespan calculated above and updates the values according to the stoichiometry of that equation. The Gillespie SSA is stochastic both with respect to the timestep used and calculation of the first reaction to fire, as shown in **Equation 3**. Because the first reaction to fire is found via cumulatively summing the probabilities of each reaction, Gillespie is somewhat dependent on the order in which reactions are defined.

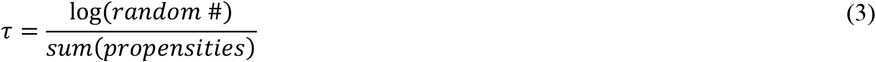

***and***

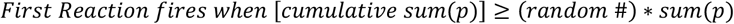

Gillespie simulation requires a defined stoichiometric matrix for each individual reaction and a vector of propensities that determine the probability of each reaction firing. Based on the system of differential equations shown in **Equation 2** and the rate constants displayed in **Table 1**, the Gillespie formulation is shown in **Equation 4**. When a reaction is determined to “fire”, the counts for each species (i.e. *tem, gen, struct*) are updated according to that reaction’s stoichiometry only.

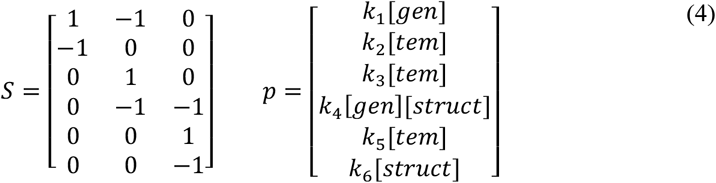

Gillespie simulation was used to generate 25 stochastic viral propagation trajectories, each with a randomly selected initial *tem* concentration. We chose to simulate very few replicates to emulate the reality of performing -omics experimentation. These varying initial conditions represent a varying severity of initial infection. Two initial *tem* concentrations were used, the first varying between 6 and 10, and the second between 4 and 14. Gillespie simulation will produce different trajectories depending on the part of the range the initial value is selected from, with the larger range producing a greater variance in trajectory. These 25 replicates are used to train an LSTM, with hyperparameters as shown in **Table 2**, and training *x* and *y* values as shown in **Equation 5**. Training *x* values include one row representing the time points for each training *y* value, with three rows which repeat the initial *x* conditions for each entry. The training *y* values are simply the simulated counts of *tem, gen*, and *struct* at each time point.

**Table 2.** Hyperparameter values used to train the viral propagation LSTM neural network.

| Hyperparameter | Value |
| --- | --- |
| # Hidden Units | 10 |
| Dropout Probability | 10% |
| Maximum Epochs | 1500 |
| Initial Learning Rate | 0.0005 |
| Fully Connected Layers | 2 (before dropout) |
|  | 3 (after dropout) |

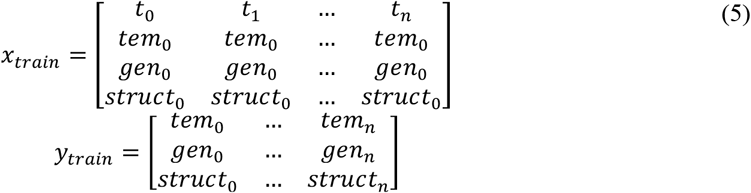

After training the LSTM network, validation was performed in multiple ways. First, the root mean squared error (RMSE) of the LSTM’s prediction of the average initial *tem* concentration (8 *tem* molecules) was calculated. The numerical solution of the viral propagation model was calculated given 8 initial *tem* molecules were used as a point of comparison. In addition, we extended this analysis by applying the elbow rule to determine the lowest number of replicates required for the LSTM to achieve acceptable accuracy. 25 LSTM networks were trained on five random subsets of data selected without replacement. These datasets contain begin containing a single replicate and one addition replicate is added for each subsequent iteration, until all 25 replicates were used. We used this technique to identify the point of diminishing returns, when adding more replicates is not a worthwhile investment. Each of these LSTM networks uses the same hyperparameters values as shown in **Table 2**.

### Aspergillus nidulans brlA Case Study

#### Strains and Media

Aspergillus nidulans A1405 (Fungal Genetics Stock Center; FGSC) was used as the control strain. Frozen stocks were spread on MAGV plates (2% malt extract, 1.5% agar, 2% glucose, 2% peptone, and 1ml/L Hutner’s trace elements and vitamin solution) and incubated for 2 days at 28 °C (29). 1E7 spores were harvested and inoculated into 50 ml of YGV (pH 3.3) (0.5% yeast extract, 1% glucose, and 1 ml/L Hutner’s trace elements and vitamin solution). Seed cultures were grown in a 250 ml baffled flask at 250 rpm and 28 °C. After 12 h growth, this flask was used to seed 1.2L YGV in a 2.8L Fernbach flask. All strains used in this study are all available from FGSC.

#### Micafungin Treatment and Extraction

After 20 h of growth (mid-exponential growth phase as determined by growth curve), various quantities of micafungin were added to the 2.8L Fernbach flask. The critical concentration of micafungin was previously determined to be 7 ng/mL in Chelius *et al*. (2020). To sample above and below this value, cultures were grown at 0, 5, 10, 15, and 20 ng/mL. 25 ml of culture from the shake flask was extracted after 0, 10, 20, 30, 60, and 90 minutes after micafungin exposure. Fungal biomass was recovered from the liquid via filtration through cheesecloth, squeezed dry, and then transferred into 25 mL centrifuge tubes by spatula. Biomass in the centrifuge tubes were flash frozen by liquid nitrogen and placed in a liquid nitrogen bath while samples were collected. Samples were placed in -80°C for at least one hour to ensure even freezing and brittleness. Frozen mycelia were crushed with mortar and pestle into a fine powder. RNA extraction, purification, and cDNA conversion were completed as described by Chelius *et al*. (2019). Based on the results of our *in silico* proof of concept using the viral model, five biological replicates and three technical replicates were collected at each time point. Each replicate’s converted cDNA was run using a BioRad C1000 Touch Thermal Cycler with CFX96Real-Time System. The target transcript *brlA* was quantified with respect to the reference gene, histone (H2B)’s transcript, and fold change was determined with respect to the zero-time point following the Ct method (52). Primers were designed using Benchling and are included in

## Supplementary Material

### LSTM Structure and Parameters

A similar training procedure was used for the *brlA* LSTM and the viral propagation one. An LSTM neural network was trained on the *brlA* transcriptomic data which takes micafungin concentration as input and predicts the dynamic response of *brlA*. **Table 3** displays the LSTM hyperparameters, which are the same for both of our *brlA* LSTM models.

**Table 3.** Hyperparameter values used to train the *brlA* transcriptomics LSTM neural network.

| Hyperparameter | Value |
| --- | --- |
| # Hidden Units | 10 |
| Dropout Probability | 10% |
| Maximum Epochs | 750 |
| Initial Learning Rate | 0.00075 |
| Fully Connected Layers | 10 (before dropout) |
|  | 3 (after dropout) |

Biological data is extremely noisy, so validation is paramount to assess the accuracy of our trained LSTMs. First for the dynamic trajectories, 75 dynamic trajectories were available among five biological replicates, three technical replicates, and five micafungin concentrations. Five-fold cross-validation was used to stress-test the network. Removing large quantities of data (15 points in each fold) was expected to lead to breakdowns in performance. For our *in vitro* case study, we selected five biological replicates based on our application of the elbow method^44^ to our *in-silico* proof of concept. Links to all code and data used to train each LSTM is included in the **Supplementary Material**.

### Surface Plotting

Once the network predicting dynamic trajectories was validated, it was used to generate *in silico* transcriptomic replicates within the range of micafungin concentration from which the dataset was generated. To do so, the network was queried from 0 to 20 ng/mL micafungin concentration in 1 ng/mL intervals. These 21 replicates were used to generate a surface plot whose *z*-axis represents *brlA*’s transcriptomic response with respect to both micafungin concentration and time after exposure. In addition to visualization via surface plot, these additional replicates could be used alongside other modeling techniques. For example, these additional replicates could be used to help fit parameters within a kinetic model, or even *de novo* generate a model via a technique such as genetic programming.

## Results and Discussion

### Viral Propagation

Despite introducing stochasticity and providing very scarce viral propagation data for training, an LSTM can accurately predict trajectories within the range of initial concentrations used for training. In addition, the LSTM shows reasonable extrapolation capability outside of the training range of initial concentrations. For the viral propagation data, an LSTM was trained on two stochastically generated datasets, one with low variation of the initial *tem* concentration, and one significantly more variation. Each LSTM was tested by generating analytical solutions for *tem* concentrations within the training range and without and calculating the error and correlation coefficient (R^2^) between analytical and LSTM-predict solutions. **Table 3** displays this analysis for both the small initial condition range (6-10 *tem* molecules) and large (4-14 *tem* molecules). **Figures 2 and 3** display representative trajectories from the small and large initial condition ranges, respectively, with each considering two close extrapolations and an interpolation.

**Figure 2.**
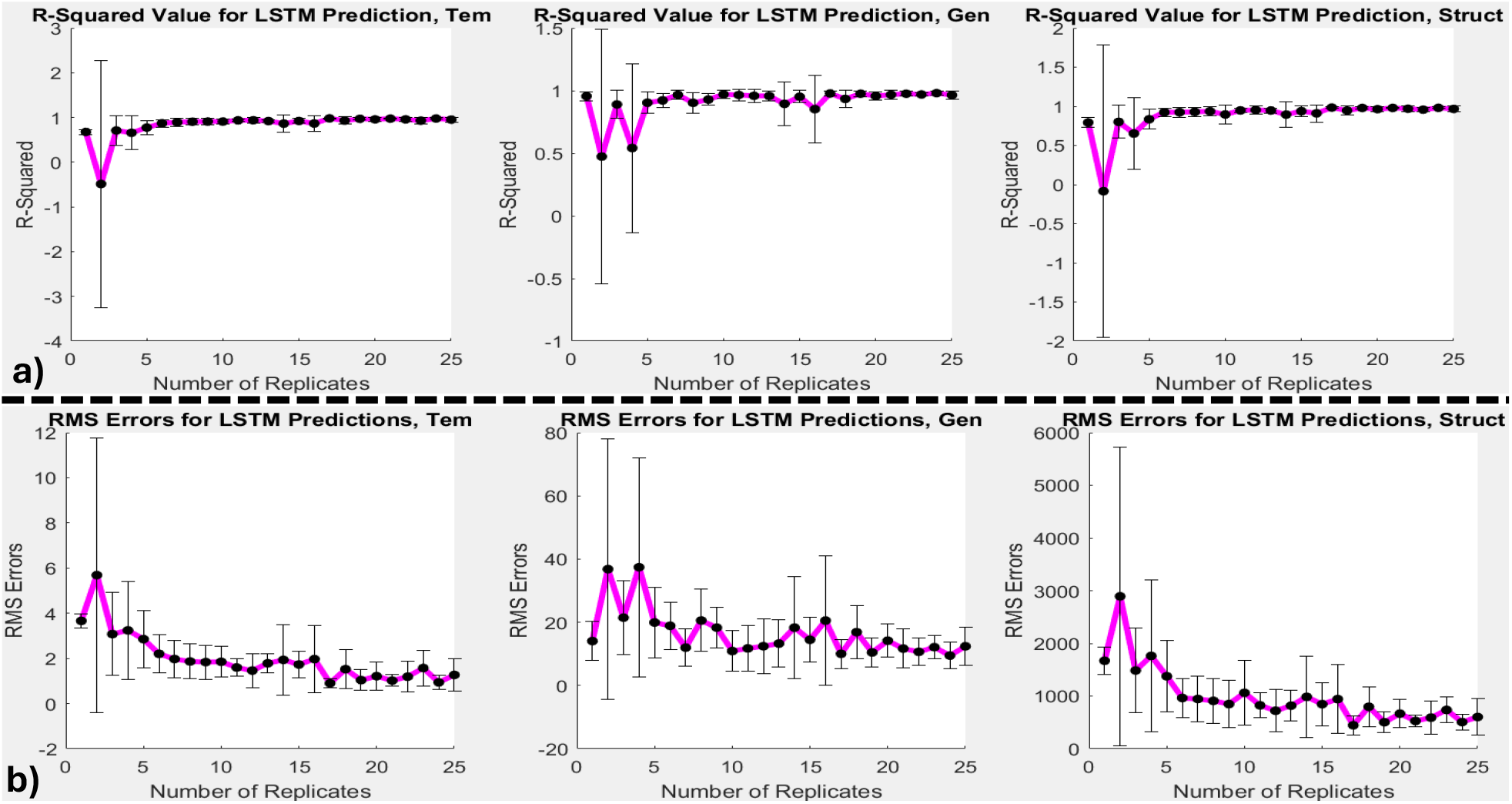
Given a small initial range of possible template (*tem*) molecules, between 6 and 10, an LSTM can predict the infection’s dynamics very accurately. To assess the accuracy of the LSTM and choose an appropriate number of replicates for our *in vitro* case study, subsets of the full dataset were selected, containing between 1 replicate and all data. For each number of replicates, five random subsets were selected without replacement and used to train an LSTM neural network. A time course at the center value of the range, 8 *tem* molecules, was generated analytically and via the LSTM, and the accuracy is reported via: **a)** R^2^, and **b)** root-mean-squared errors for *tem, gen*, and *struct*. We suggest that an elbow occurs at five replicates, indicating that additional replicates beyond this point provide diminishing return.

**Figure 3.**
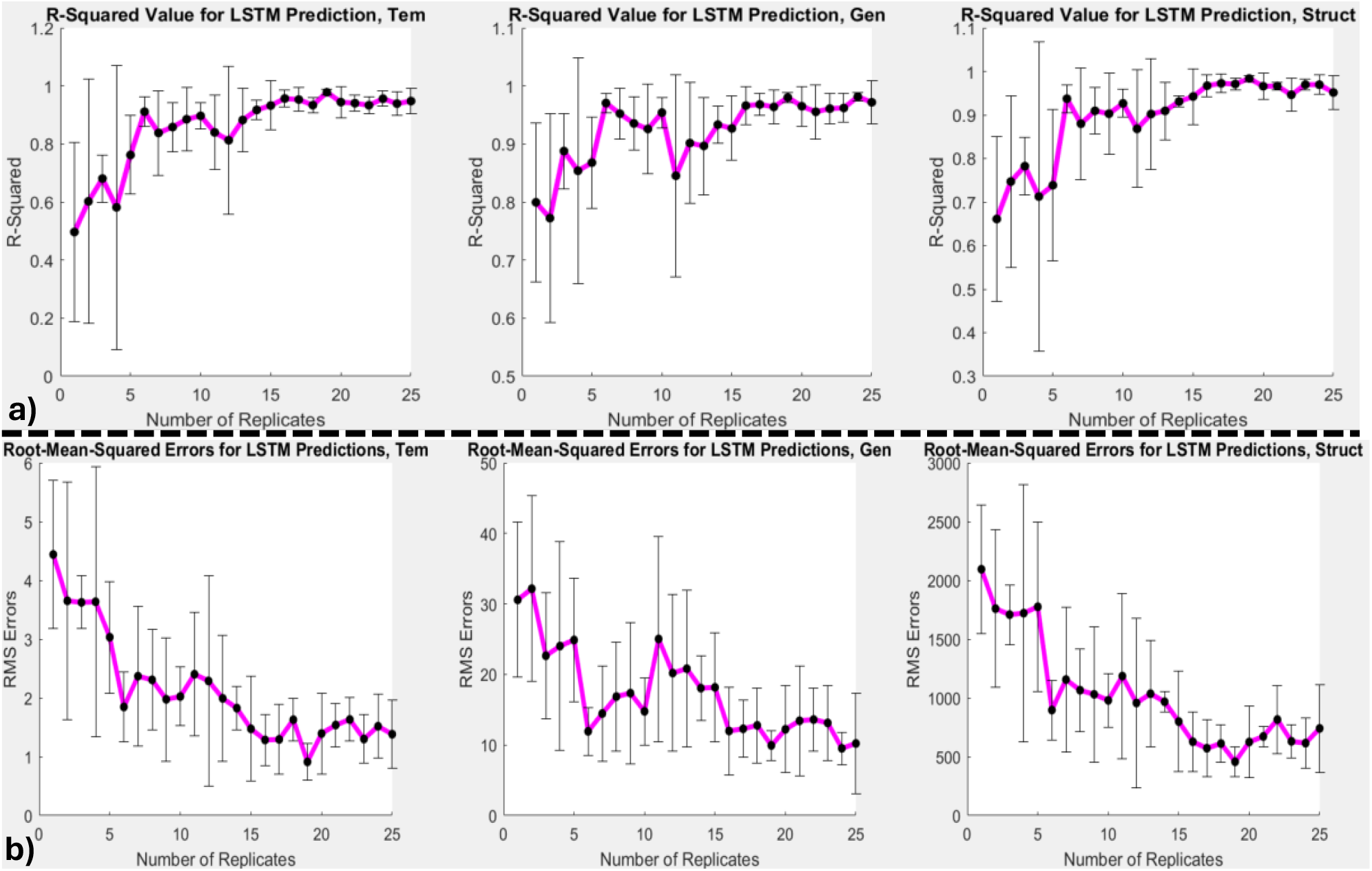
Given a larger initial range of possible template (*tem*) molecules, between 4 and 12, an LSTM can predict the dynamics of infection accurately but requires more replicates to do so. This exercise is likely more realistic to the amount of noise present in biological data, and what can be expected in the *in vitro* case study. As in **Figure 2**, a time course at the center value of the range, 8 *tem* molecules, was generated analytically and via the LSTM, and the accuracy is reported via: **a)** R^2^, and **b)** root-mean-squared errors for *tem, gen*, and *struct*. We suggest that an elbow occurs at six replicates, indicating that additional replicates beyond this point provide diminishing return.

For the small initial condition range, LSTM shows good performance for all interpolation cases, including those at the edge of the possible initial conditions. However, the LSTM shows significantly worse performance extrapolating solutions with initial conditions greater than the training dataset, with smaller initial conditions retaining high R^2^ values. A possible cause for this imbalance is that the randomly selected initial *tem* values in the training dataset were skewed toward small values. However, with 25 replicates, it is highly unlikely that such a large skew exists. A more likely cause is that higher initial viral loads lead to different infection and propagation dynamics, confusing the LSTM which has not been exposed to those dynamics. Compare the extrapolated trajectories in **Figure 2(b) and (c)**, in which high initial *tem* concentrations lead to significantly less lag time in both *gen* and *struct*, as well as a faster and more intense initial dip in *tem* concentration.

For the large initial condition range, the LSTM has lower R^2^ values and worse RMSE for all trajectories. This result is expected, especially due to the known significant difference in dynamics between severe and mild infections. In addition, the large initial *tem* range has a more pronounced asymmetry with regard to validation R^2^ and RMSE at low and high initial conditions, as shown in **Table 4**. The extrapolation ability of the LSTM is diminished in the larger initial *tem* range. This effect is expected both due to using a wider range with the same number of replicates, and the fact that expanding the extremes introduces more varied dynamics.

**Table 4.** An error assessment for interpolated and extrapolated time-courses for LSTMs trained on datasets generated with small (6-10) and large (4-12) initial template concentration ranges. Italicized values represent extrapolations, bold values represent the center value of the initial condition range. The final section reports the error for the large (4-12) initial range but seeded with five extra replicates in the top half (8-12) of the range.

| 6-10 initial template |  |  |  |  |  |  |
| --- | --- | --- | --- | --- | --- | --- |
| $tem_0$ | Rsq,T | Rsq,G | Rsq,S | RMSE,T | RMSE,G | RMSE,S |
| 4 | 0.9784 | 0.9884 | 0.9537 | 1.1103 | 8.4509 | 861.507 |
| 6 | 0.9811 | 0.9962 | 0.9815 | 0.9604 | 4.6157 | 522.25 |
| **8** | **0.9565** | **0.9732** | **0.9751** | **1.3472** | **11.7476** | **582.903** |
| 10 | 0.8152 | 0.8727 | 0.8776 | 2.5755 | 24.742 | 1251.3 |
| 11 | 0.6242 | 0.7538 | 0.7496 | 3.5411 | 33.8824 | 1761.1 |
| 12 | 0.241 | 0.5494 | 0.5147 | 4.8568 | 45.1871 | 2415.7 |
| 4-12 initial template |  |  |  |  |  |  |
| $tem_0$ | Rsq,T | Rsq,G | Rsq,S | RMSE,T | RMSE,G | RMSE,S |
| 3 | 0.4657 | 0.5312 | 0.4434 | 5.7425 | 55.093 | 3059 |
| 4 | 0.6829 | 0.7374 | 0.6769 | 4.2528 | 40.1312 | 2276.8 |
| 6 | 0.9405 | 0.9574 | 0.956 | 1.7022 | 15.6762 | 792.126 |
| **8** | **0.8813** | **0.899** | **0.9359** | **2.225** | **22.7995** | **936.102** |
| 10 | 0.5673 | 0.6986 | 0.7389 | 3.9404 | 38.0636 | 1827.6 |
| 12 | -0.0159 | 0.4285 | 0.412 | 5.6189 | 51.6195 | 2621.3 |
| 13 | -0.4607 | 0.2305 | 0.2178 | 6.511 | 58.2656 | 3024.3 |
| 4-12 initial template, seeded |  |  |  |  |  |  |
| $tem_0$ | Rsq,T | Rsq,G | Rsq,S | RMSE,T | RMSE,G | RMSE,S |
| 3 | 0.9018 | 0.9077 | 0.9157 | 2.4617 | 24.4433 | 1190.6 |
| 4 | 0.9455 | 0.9552 | 0.9551 | 1.7637 | 16.5681 | 848.7297 |
| 6 | 0.9819 | 0.9935 | 0.9858 | 0.9378 | 6.0434 | 457.4024 |
| **8** | **0.9486** | **0.9718** | **0.9693** | **1.4636** | **12.0571** | **647.4241** |
| 10 | 0.8091 | 0.8871 | 0.8955 | 2.6177 | 23.2943 | 1156.2 |
| 12 | 0.5448 | 0.7515 | 0.7683 | 3.7612 | 33.5555 | 1669.3 |
| 13 | 0.374 | 0.6777 | 0.6958 | 4.2624 | 37.7103 | 1885.9 |

To address the asymmetry in error, we seeded the dataset with five additional stochastically generated replicates with initial conditions in the top half of the range. Seeding additional replicates within this problem region drastically improved its predictive performance through the entire range of initial conditions. In fact, as seen in **Table 4**, just five additional replicates drastically improve error at the extremes of the initial condition range. The LSTM still struggles to predict the complex dynamics at high initial conditions, but additional steps such as applying weights to these initial conditions during training could further improve accuracy. Most interestingly, adding five additional high initial *tem* replicates drastically improved and extended the accuracy of the LSTM when predicting low initial *tem* replicates. This effect suggests that addressing the weakest link can benefit prediction among the entire dataset.

We applied the elbow rule to both initial condition ranges both to further validate the LSTMs and also the number of replicates required to reach satisfactory performance.^44^ The purpose of the elbow rule is to identify the “point of diminishing returns,” which is essentially the minimum number of biological replicates required to get acceptable predictive performance should this type of experiment be performed *in vitro*. **Figures 4 and 5** display elbow rule visualizations for both the small and large initial condition ranges, respectively. We have identified the “elbow” or point of diminishing returns for each range to be five and six replicates, respectively. The error metrics decrease in variance and improve at a significantly slower rate beyond five and six replicates, respectively. While the elbow rule is largely qualitative, it is a valuable first pass to help decide how many replicates are necessary for accurate *in vitro* experimentation and modeling. Based on the results of this proof of concept, we decided to collect five biological replicates for our *in vitro* case study using *Aspergillus nidulans* BrlA.

**Figure 4.**
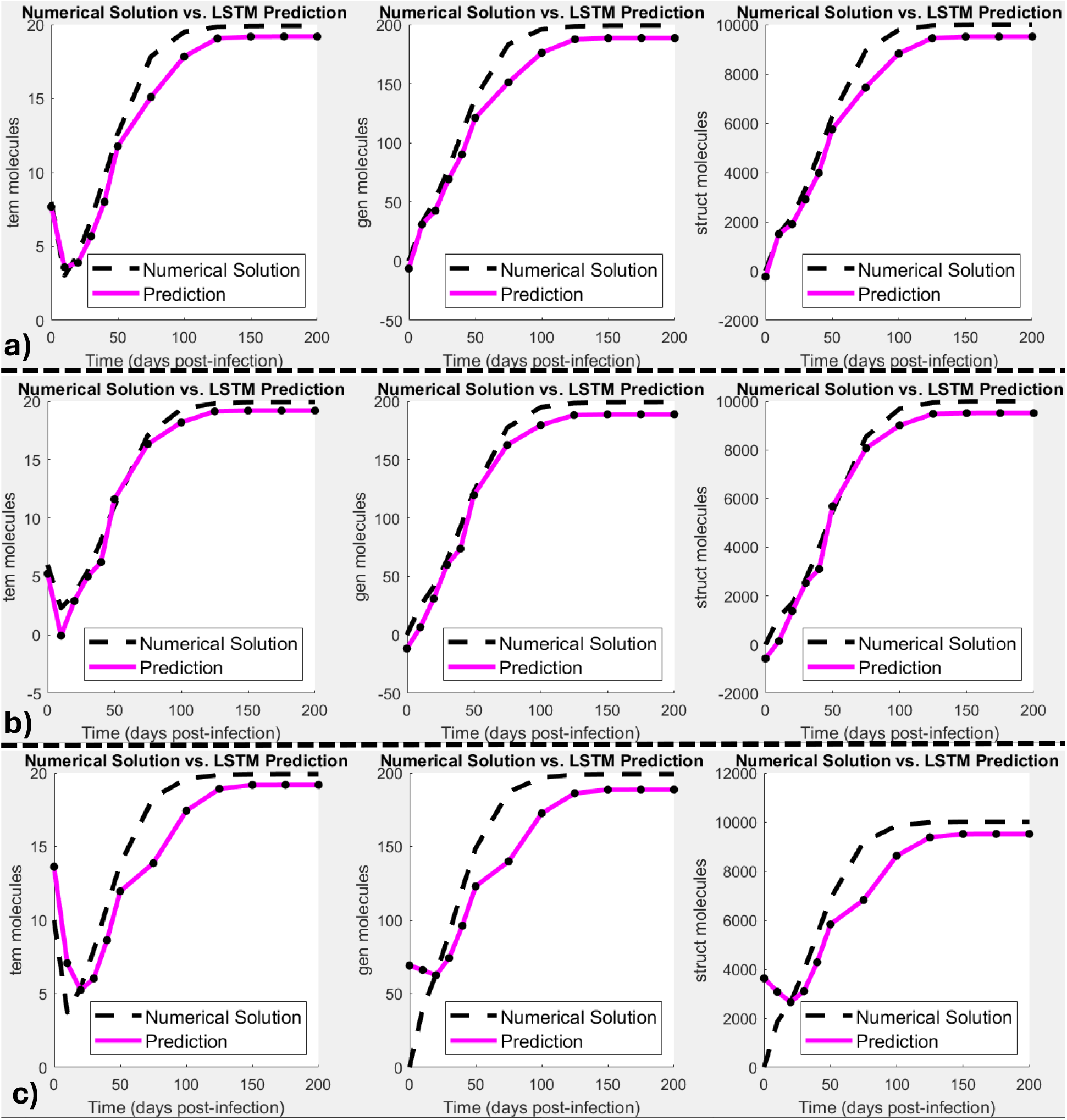
Given a smaller initial range of possible template (*tem*) molecules, between 6 and 10, an LSTM can predict the dynamics of infection accurately. Each row represents LSTM time course predictions for *tem, gen*, and *struct* at initial *tem* values of **a)** 6 *tem* molecules, **b)** 8 *tem* molecules, **c)** 10 *tem* molecules. The predictions at 6 and 10 are just at the edge of the available training range. Near the bottom of the training dataset range, prediction remains highly accurate, while predictions near the top of the range result in significant errors. This is likely because infections with high initial *tem* concentration result in more extreme dynamics early in the trajectory, which differs from lower initial concentrations.

**Figure 5.**
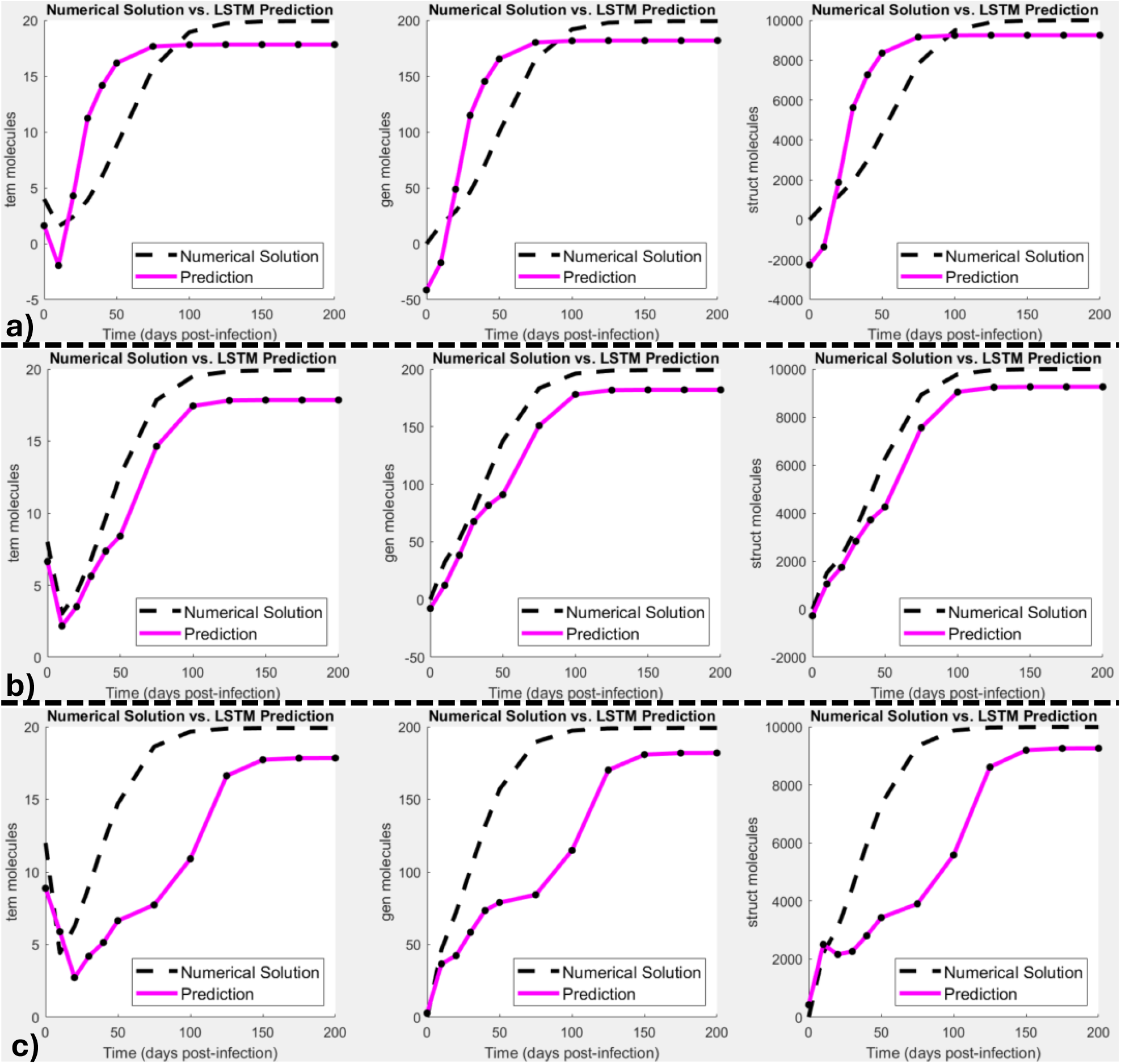
Given a larger initial range of possible template (*tem*) molecules, between 4 and 12, an LSTM is less accurate in predicting propagation time courses. Each row represents LSTM time course predictions for *tem, gen*, and *struct* at initial *tem* values of **a)** 4 *tem* molecules, **b)** 8 *tem* molecules, **c)** 12 *tem* molecules. The predictions at 4 and 12 are just at the edge of the available training range. Near the bottom of the training dataset range, prediction remains more accurate than top of the range, as in **Figure 4**. By seeding the neural network with five additional replicates in the top of the range, shown in **d)**, the prediction at 12 *tem* molecules drastically improves.

### Aspergillus nidulans brlA Case Study

For noisy biological data with a wide range of micafungin concentrations, the LSTM’s validation R^2^ value is acceptable and in line with expected accuracy based on our viral propagation proof of concept. In five-fold cross-validation, the average R^2^ achieved by the dynamic trajectory LSTM is approximately 0.71, as displayed in **Table 5**. The relatively low R^2^ value is likely due to the inconsistent outlier nature of the data at 10 ng/mL, discussed later. This behavior confounds our LSTM’s prediction capability, especially when replicates that would reinforce this paradoxical behavior are left out during validation. These results at the point of greatest paradoxical behavior are still a drastic improvement over linear interpolation. Interpolation techniques would likely completely miss this peak, but LSTMs trained on full time-course trajectories can somewhat capture the true dynamics.

The main focus of this paper is on the development of an LSTM technique for modeling dynamic -omics trajectories and extracting additional data for use in model development. However, our transcriptomic data also reveal a paradoxical trend in *brlA* transcriptional regulation in response to micafungin exposure. As a point of clarification, there is already a well-known “paradoxical effect” between *Aspergilli* and echinocandin antifungals.^45^ *Aspergilli* paradoxically recover the ability to grow effectively at high echinocandin concentrations, which has been shown in many echinocandins.^46–49^ The paradoxical effect we will reference for the rest of the paper is transcriptomic, not related directly to growth rate; however, these two effects may be linked. The paradoxical transcriptomic response of *brlA* is further exemplified by the surface plot generated from additional *in-silico* replicates from our LSTM. The expected dose-response behavior is that higher doses elicit a higher response, unless the dose is lethal.

Based on our previously published data ^39,40^, no concentration within our range of micafungin doses is lethal. We generated a surface plot to visualize this paradoxical effect by querying our LSTM with many micafungin concentrations as input, which is shown in **Figure 6**. The surface plot illustrates the paradoxical behavior of *A. nidulans* BrlA, in which 10 ng/mL micafungin elicits the greatest transcriptional response.

**Figure 6.**
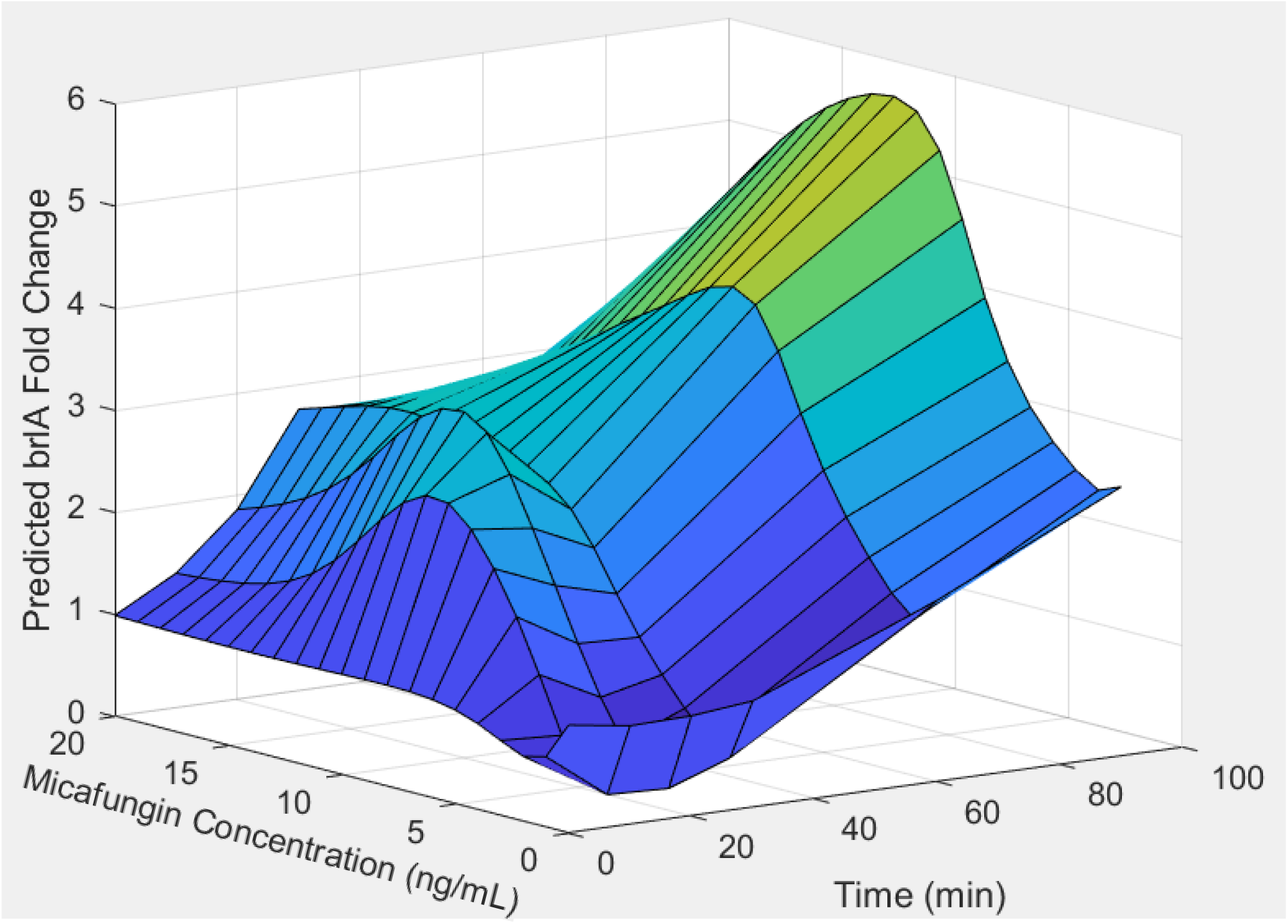
An LSTM trained on all *brlA* transcriptomic data was used to generate the following surface plot, by querying at 1 ng/mL intervals from 0 to 20 ng/mL of micafungin. This surface plot illustrates micafungin’s “paradoxical effect” on *brlA* expression. Medium doses of micafungin elicit the strongest *brlA* response, while higher doses cause a weaker response. In addition, medium micafungin concentrations elicit an initial spike in *brlA* transcription, followed by the more typical exponential rise seen in all other concentrations. In previous work, Chelius *et al*. (2020), the lowest concentration at which micafungin caused visual impairment was between 7 and 10 ng/mL. It is possible that the fungistatic effect of micafungin kicks in very quickly and hinders the fungi, or that the *brlA* response is only effective at low micafungin concentrations.

Additionally, all dynamic trajectories have a steady, exponential increase in fold change with respect to time after micafungin exposure. Again, 10 ng/mL appears to have the highest average transcription response at each time point. However, at 20 minutes after exposure, 10 ng/mL micafungin elicits a secondary spike in *brlA* transcript which then quickly fades as shown in **Figure 6**. Both the overall dose-response behavior and the dynamic transcriptomic trajectory of 10 ng/mL display paradoxical and/or counterintuitive responses. Namely, 10 ng/mL is consistently at higher regulation levels, and shows a unique, intense spike at 20 minutes.

Finally, we wish to address our use of the elbow method to inform usage of five biological replicates. Within scientific literature, there is no concrete measure for how many biological replicates are enough to capture a biological effect and accurately draw conclusions. Many researchers operate with the rule of thumb of two to three biological replicates and three technical replicates. Due to high variance, we found that applying this machine learning technique to -omics data requires at least five biological replicates. We will not claim that our treatment is a concrete rule for all biological research, but we do suggest that researchers following the common rule of thumb may not yet have reached the point of diminishing returns with regards to the number of replicates used. Researchers collecting only two or three biological replicates may be losing valuable information about the systems they are studying.

## Conclusion

The long short-term memory neural network is an effective method to capture and amplify trends within noisy and/or sparse biological data. We have successfully conducted an *in silico* proof of concept and an *in vitro* case study, both of which demonstrate an LSTM’s ability to generate additional *in silico* time-resolved replicates for -omics studies. Synthetic -omics time courses can be generated on-demand by querying the trained neural network. Both the biological and synthetic time courses can be used to generate mechanistic models, such as in genetic programming. A key barrier to the effective use of machine learning in biology is the cost and time investment of producing a sufficient quantity of replicates. We have demonstrated that LSTM neural networks can produce additional *in silico* replicates for use in machine learning techniques that require large datasets.

## Supporting information

Supplemental Table and Links

## Acknowledgements

This material is based upon work supported by the National Science Foundation under Grant No. 2006190 and No. 2006189.

## References

(1) Crick, F. Central Dogma of Molecular Biology. Nature 1970, 227 (5258), 561–563. 10.1038/227561a0.

(2) Zhao, Z.; Meng, F.; Wang, W.; Wang, Z.; Zhang, C.; Jiang, T. Comprehensive RNA-Seq Transcriptomic Profiling in the Malignant Progression of Gliomas. Sci Data 2017, 4 (1), 170024. 10.1038/sdata.2017.24.

(3) Kim, H.-J.; Rakwal, R.; Shibato, J.; Iwahashi, H.; Choi, J.-S.; Kim, D.-H. Effect of Textile Wastewaters on Saccharomyces Cerevisiae Using DNA Microarray as a Tool for Genome-Wide Transcriptomics Analysis. Water Res 2006, 40 (9), 1773–1782. 10.1016/j.watres.2006.02.037.

(4) Giménez, M. J.; Pistón, F.; Atienza, S. G. Identification of Suitable Reference Genes for Normalization of QPCR Data in Comparative Transcriptomics Analyses in the Triticeae. Planta 2011, 233 (1), 163–173. 10.1007/s00425-010-1290-y.

(5) Aebersold, R.; Mann, M. Mass Spectrometry-Based Proteomics. Nature 2003, 422 (6928), 198–207. 10.1038/nature01511.

(6) Nørregaard Jensen, O. Modification-Specific Proteomics: Characterization of Post-Translational Modifications by Mass Spectrometry. Curr Opin Chem Biol 2004, 8 (1), 33– 41. 10.1016/j.cbpa.2003.12.009.

(7) Slobodyanyuk, M.; Bahcheli, A. T.; Klein, Z. P.; Bayati, M.; Strug, L. J.; Reimand, J. Directional Integration and Pathway Enrichment Analysis for Multi-Omics Data. Nat Commun 2024, 15 (1), 5690. 10.1038/s41467-024-49986-4.

(8) Reimand, J.; Isserlin, R.; Voisin, V.; Kucera, M.; Tannus-Lopes, C.; Rostamianfar, A.; Wadi, L.; Meyer, M.; Wong, J.; Xu, C.; Merico, D.; Bader, G. D. Pathway Enrichment Analysis and Visualization of Omics Data Using g:Profiler, GSEA, Cytoscape and EnrichmentMap. Nat Protoc 2019, 14 (2), 482–517. 10.1038/s41596-018-0103-9.

(9) Ahmed, S.; Zhang, M.; Peng, L. Genetic Programming for Biomarker Detection in Mass Spectrometry Data; 2012; pp 266–278. 10.1007/978-3-642-35101-3_23.

(10) Ahmed, S.; Zhang, M.; Peng, L. Improving Feature Ranking for Biomarker Discovery in Proteomics Mass Spectrometry Data Using Genetic Programming. Conn Sci 2014, 26 (3), 215–243. 10.1080/09540091.2014.906388.

(11) Hu, T.; Oksanen, K.; Zhang, W.; Randell, E.; Furey, A.; Zhai, G. Analyzing Feature Importance for Metabolomics Using Genetic Programming; 2018; pp 68–83. 10.1007/978-3-319-77553-1_5.

(12) Murray, P. G.; Stevens, A.; De Leonibus, C.; Koledova, E.; Chatelain, P.; Clayton, P. E. Transcriptomics and Machine Learning Predict Diagnosis and Severity of Growth Hormone Deficiency. JCI Insight 2018, 3 (7). 10.1172/jci.insight.93247.

(13) Swan, A. L.; Mobasheri, A.; Allaway, D.; Liddell, S.; Bacardit, J. Application of Machine Learning to Proteomics Data: Classification and Biomarker Identification in Postgenomics Biology. OMICS 2013, 17 (12), 595–610. 10.1089/omi.2013.0017.

(14) Blainey, P.; Krzywinski, M.; Altman, N. Replication. Nat Methods 2014, 11 (9), 879–880. 10.1038/nmeth.3091.

(15) Hochreiter, S.; Schmidhuber, J. Long Short-Term Memory. Neural Comput 1997, 9 (8), 1735–1780. 10.1162/neco.1997.9.8.1735.

(16) Denning, D. W. Echinocandin Antifungal Drugs. The Lancet 2003, 362 (9390), 1142– 1151. 10.1016/S0140-6736(03)14472-8.

(17) Grover, N. Echinocandins: A Ray of Hope in Antifungal Drug Therapy. Indian J Pharmacol 2010, 42 (1), 9. 10.4103/0253-7613.62396.

(18) Srivastava, R.; You, L.; Summers, J.; Yin, J. Stochastic vs. Deterministic Modeling of Intracellular Viral Kinetics. J Theor Biol 2002, 218 (3), 309–321. 10.1006/jtbi.2002.3078.

(19) Yao, K.; Zweig, G.; Hwang, M.-Y.; Shi, Y.; Yu, D. Recurrent Neural Networks for Language Understanding. In Interspeech 2013; ISCA: ISCA, 2013; pp 2524–2528. 10.21437/Interspeech.2013-569.

(20) Sherstinsky, A. Fundamentals of Recurrent Neural Network (RNN) and Long Short-Term Memory (LSTM) Network. Physica D 2020, 404, 132306. 10.1016/j.physd.2019.132306.

(21) Sherstinsky, A. Deriving the Recurrent Neural Network Definition and RNN Unrolling Using Signal Processing. In Critiquing and Correcting Trends in Machine Learning Workshop (CRACT) -NeurIPS 2018; 2018.

(22) Bengio, Y.; Frasconi, P.; Simard, P. The Problem of Learning Long-Term Dependencies in Recurrent Networks. In IEEE International Conference on Neural Networks; IEEE; pp 1183–1188. 10.1109/ICNN.1993.298725.

(23) Hochreiter, S. The Vanishing Gradient Problem During Learning Recurrent Neural Nets and Problem Solutions. International Journal of Uncertainty, Fuzziness and Knowledge-Based Systems 1998, 06 (02), 107–116. 10.1142/S0218488598000094.

(24) Basodi, S.; Ji, C.; Zhang, H.; Pan, Y. Gradient Amplification: An Efficient Way to Train Deep Neural Networks. Big Data Mining and Analytics 2020, 3 (3), 196–207. 10.26599/BDMA.2020.9020004.

(25) Nair, V.; Hinton, G. E. Rectified Linear Units Improve Restricted Boltzmann Machines. In Proceedings of the 27th International Conference on Machine Learning; 2010.

(26) Gers, F. A.; Schraudolph, N. N.; Schmidhuber, J. Learning Precise Timing with LSTM Recurrent Networks. Journal of Machine Learning Research 2002, 3, 115–143.

(27) Gers, F. A.; Schmidhuber, J.; Cummins, F. Learning to Forget: Continual Prediction with LSTM. Neural Comput 2000, 12 (10), 2451–2471. 10.1162/089976600300015015.

(28) Li, C.; Zhou, J.; Du, G.; Chen, J.; Takahashi, S.; Liu, S. Developing Aspergillus Niger as a Cell Factory for Food Enzyme Production. Biotechnol Adv 2020, 44, 107630. 10.1016/j.biotechadv.2020.107630.

(29) Hu, W.; Liu, Z.; Fu, B.; Zhang, X.; Qi, Y.; Hu, Y.; Wang, C.; Li, D.; Xu, N. Metabolites of the Soy Sauce Koji Making with Aspergillus Niger and Aspergillus Oryzae. Int J Food Sci Technol 2022, 57 (1), 301–309. 10.1111/ijfs.15406.

(30) Upton, D. J.; McQueen-Mason, S. J.; Wood, A. J. An Accurate Description of Aspergillus Niger Organic Acid Batch Fermentation through Dynamic Metabolic Modelling. Biotechnol Biofuels 2017, 10 (1), 258. 10.1186/s13068-017-0950-6.

(31) Latgé, J.-P. The Pathobiology of Aspergillus Fumigatus. Trends Microbiol 2001, 9 (8), 382–389. 10.1016/S0966-842X(01)02104-7.

(32) Kwon-Chung, K. J.; Sugui, J. A. Aspergillus Fumigatus—What Makes the Species a Ubiquitous Human Fungal Pathogen? PLoS Pathog 2013, 9 (12), e1003743. 10.1371/journal.ppat.1003743.

(33) Zhang, X.; Zhu, Y.; Bao, L.; Gao, L.; Yao, G.; Li, Y.; Yang, Z.; Li, Z.; Zhong, Y.; Li, F.; Yin, H.; Qu, Y.; Qin, Y. Putative Methyltransferase LaeA and Transcription Factor CreA Are Necessary for Proper Asexual Development and Controlling Secondary Metabolic Gene Cluster Expression. Fungal Genetics and Biology 2016, 94, 32–46. 10.1016/j.fgb.2016.07.004.

(34) Son, Y.-E.; Yu, J.-H.; Park, H.-S. Regulators of the Asexual Life Cycle of Aspergillus Nidulans. Cells 2023, 12 (11), 1544. 10.3390/cells12111544.

(35) Clutterbuck, A. J. A MUTATIONAL ANALYSIS OF CONIDIAL DEVELOPMENT IN ASPERGILLUS NIDULANS. Genetics 1969, 63 (2), 317–327. 10.1093/genetics/63.2.317.

(36) Sewall, T. C. Cellular Effects of Misscheduled BrlA, AbaA, and WetA Expression in Aspergillus Nidulans. Can J Microbiol 1994, 40 (12), 1035–1042. 10.1139/m94-164.

(37) Rocha, M. C.; Fabri, J. H. T. M.; Simões, I. T.; Silva-Rocha, R.; Hagiwara, D.; da Cunha, A. F.; Goldman, G. H.; Cánovas, D.; Malavazi, I. The Cell Wall Integrity Pathway Contributes to the Early Stages of Aspergillus Fumigatus Asexual Development. Appl Environ Microbiol 2020, 86 (7). 10.1128/AEM.02347-19.

(38) Kovács, Z.; Szarka, M.; Kovács, S.; Boczonádi, I.; Emri, T.; Abe, K.; Pócsi, I.; Pusztahelyi, T. Effect of Cell Wall Integrity Stress and RlmA Transcription Factor on Asexual Development and Autolysis in Aspergillus Nidulans. Fungal Genetics and Biology 2013, 54, 1–14. 10.1016/j.fgb.2013.02.004.

(39) Reese, S.; Chelius, C.; Riekhof, W.; Marten, M. R.; Harris, S. D. Micafungin-Induced Cell Wall Damage Stimulates Morphological Changes Consistent with Microcycle Conidiation in Aspergillus Nidulans. Journal of Fungi 2021, 7 (7), 525. 10.3390/jof7070525.

(40) Chelius, C.; Huso, W.; Reese, S.; Doan, A.; Lincoln, S.; Lawson, K.; Tran, B.; Purohit, R.; Glaros, T.; Srivastava, R.; Harris, S. D.; Marten, M. R. Dynamic Transcriptomic and Phosphoproteomic Analysis During Cell Wall Stress in Aspergillus Nidulans. Molecular & Cellular Proteomics 2020, 19 (8), 1310–1329. 10.1074/mcp.RA119.001769.

(41) Espinel-Ingroff, A. In Vitro Antifungal Activities of Anidulafungin and Micafungin, Licensed Agents and the Investigational Triazole Posaconazole as Determined by NCCLS Methods for 12,052 Fungal Isolates: Review of the Literature. Rev Iberoam Micol 2003, 20 (4), 121–136.

(42) Edwards, H.; Zavorskas, J.; Huso, W.; Doan, A. G.; Grey, K.; Lee, J.; Morse, M.; Wilkinson, H. H.; Ebbole, D.; Shaw, B. D.; Harris, S. D.; Srivastava, R.; Marten, M. R. Aspergillus Nidulans Transcription Factor BrlA Is Utilized in a Conidiation-Independent Response to Cell-Wall Stress. November 21, 2024. 10.1101/2024.11.21.624663.

(43) Gillespie, D. T. Stochastic Simulation of Chemical Kinetics. Annu Rev Phys Chem 2007, 58 (1), 35–55. 10.1146/annurev.physchem.58.032806.104637.

(44) Humaira, H.; Rasyidah, R. Determining The Appropiate Cluster Number Using Elbow Method for K-Means Algorithm. In Proceedings of the Proceedings of the 2nd Workshop on Multidisciplinary and Applications (WMA) 2018, 24-25 January 2018, Padang, Indonesia; EAI, 2020. 10.4108/eai.24-1-2018.2292388.

(45) Loiko, V.; Wagener, J. The Paradoxical Effect of Echinocandins in Aspergillus Fumigatus Relies on Recovery of the β-1,3-Glucan Synthase Fks1. Antimicrob Agents Chemother 2017, 61 (2). 10.1128/AAC.01690-16.

(46) Steinbach, W. J.; Lamoth, F.; Juvvadi, P. R. Potential Microbiological Effects of Higher Dosing of Echinocandins. Clinical Infectious Diseases 2015, 61 (Suppl_6), S669–S677. 10.1093/cid/civ725.

(47) Wiederhold, N. P. Paradoxical Echinocandin Activity: A Limited in Vitro Phenomenon? Med Mycol 2009, 47 (1), S369–S375. 10.1080/13693780802428542.

(48) Walker, L. A.; Gow, N. A. R.; Munro, C. A. Fungal Echinocandin Resistance. Fungal Genetics and Biology 2010, 47 (2), 117–126. 10.1016/j.fgb.2009.09.003.

(49) Hall, G. S.; Myles, C.; Pratt, K. J.; Washington, J. A. Cilofungin (LY121019), an Antifungal Agent with Specific Activity against Candida Albicans and Candida Tropicalis. Antimicrob Agents Chemother 1988, 32 (9), 1331–1335. 10.1128/AAC.32.9.1331.

