## Supplemental Table and Links for "Synthetic Generation of Dynamic Omics Data Demonstrates *Aspergillus nidulans* BrlA Paradoxical Wall Stress Response"

S1 – Code and Data Availability

All code for needed to reproduce LSTM results are included on GitHub at https://github.com/jzavorskas/LSTM.

All qPCR data used in the study is also available in that GitHub repository for download.

S2 – Lists of primers used in the qPCR study.

| | |
|---|---|
| BrlA-qPCR-Fwd | TCTCCTACACCAACTCCAACAA |
| BrlA-qPCR-Rvs | ATTCGTTCCTGCCCTTCC |
| histone2b-qPCR-Fwd | CACCCGGACACTGGTATCTC |
| histone2b-qPCR-Rvs | GAATACTTCGTAACGGCCTTGG |
